# Differential Gene Expression Pipeline for Whole Transcriptome RNA-Seq Data using Personal Computer

**DOI:** 10.1101/2021.01.26.428352

**Authors:** Rashid Saif, Aniqa Ejaz, Tania Mahmood, Saeeda Zia

## Abstract

Advances in the next generation sequencing (NGS) technologies, their cost effectiveness and well-developed pipelines using computational tools/softwares has allowed researchers to reveal ground-breaking discoveries in multi-omics data analysis. However, there is still uncertainty due to massive upsurge in parallel tools and difficulty in choosing best practiced pipeline for expression profiling of RNA sequenced (RNA-seq) data. Here, we detail the optimized pipeline that works at a fast pace with enhanced accuracy on personal computer rather than using cloud or high-performance computing clusters (HPC). The steps include quality check, base filtration, quasi-mapping, quantification of samples, estimation and counting of transcript/gene expression abundances, identification and clustering of differentially expressed features and visualization of the data. The tools FastQC, Trimmomatic, Salmon and some other scripts in Trinity toolkit were applied on two paired-end datasets. An extension of this pipeline may also be formulated in future for the gene ontology enrichment analysis and functional annotation of the differential expression matrix to make this data biologically more significant.

## INTRODUCTION

High throughput sequencing technology with rapid RNA-Seq efforts along with reduced cost has enabled the advent of powerful NGS data analysis. It has become a standard method and has superseded hybridization-based technologies such as microarrays in detailed profiling and quantification of gene and RNA expression levels respectively (Rapaport et al. 2013). The great potential of RNA-Seq to differential gene expression (DGE) technology along with its vast dynamic range have popped unrivalled progress in transcriptomics research. It also urged the computational biologists to develop and update several computational tools based on mathematical algorithms for analyzing the large data promptly (Finotello and Di Camillo 2015).

The preferred protocol for RNA-Seq experiment starts with the fragmentation of RNAs in a sample of interest and reverse-transcribing all RNAs into cDNA. Fragments are subjected to NGS which generates millions of short reads that can be further mapped on to reference genome (Yalamanchili et al. 2017). The number of reads aligned to each gene or transcript (counts) represents the digital measure of gene or transcript expression levels in the subject sample (Van Verk et al. 2013). These summarized data are affected by factors such as gene length, library size, and sequencing depth biases. These artifacts must be addressed by normalization methods that leads to list of genes with P-values, fold changes (FC) and counts per million reads (cpm) (Conesa et al. 2016). The measures Reads Per Kilobase of transcript per Million mapped reads (RPKM), Fragments Per Kilobase of transcript per Million mapped reads (FPKM), transcript per million (TPM), Trimmed Means of M values (TMM) etc. also reports normalized RNA-seq gene expression values (Qi et al. 2017). Several sophisticated algorithms and tools for differential expression have been developed viz. salmon, sailfish, kallisto, HTSeq, edgeR, DESeq, baySeq, voom, trinity toolkit and others (Robinson et al. 2010; Patro et al. 2017).

In this report, we propose an efficient and vigorous protocol focusing on each computational step, from quality assessment of RNA-Seq data to DGE analysis, normalization of raw read counts and its visualization from an exemplary dataset containing 4 raw sequence files (pair from each organism). The scheme for RNA-Seq data analysis depicted in this paper will provide a map to other users to implement or analyze their own data seamlessly.

## MATERIALS

### Equipment

1. Ubuntu v 20.04 https://ubuntu.com/
2. R and RStudio software https://cran.r-project.org/bin/windows/base/
3. FASTQC https://sourceforge.net/projects/fastqc.mirror/
4. Trimmomatic v. 0.36 https://astrobiomike.github.io/bash/installing_tools#trimmomatic
5. Salmon https://github.com/COMBINE-lab/salmon/archive/master.zip
6. Trinity https://github.com/trinityrnaseq/trinityrnaseq/archive/master.zip
7. edgeR, limma, DESeq2, ctc, Biobase on R. https://cran.rproject.org/web/packages/BiocManager/index.html

## METHOD

### Protocol for DGE analysis of RNA-Seq data

The raw fastq files is subjected to different steps as depicted in (Fig. 1).

**Figure 1.**
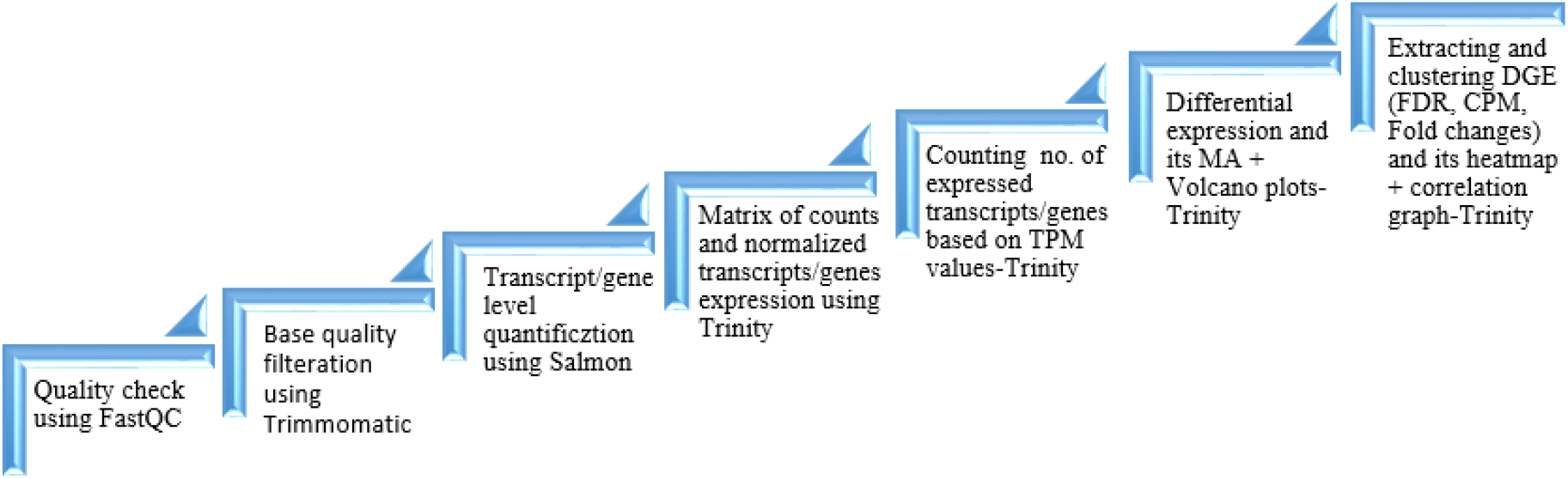
Workflow for RNA-Seq data. The pipeline for differential transcript/gene expression analysis of RNA-seq data

The pipeline is an integration of various tools viz. FastQC, Trimmomatic, Salmon and Trinity toolkit. The pipeline was run on Linux (Ubuntu v. 20.04 LTS) operating system with 2.60 GHz processor, 2 core(s) CPU having 8 GB RAM.

### Quality checks and base quality filtration

**1.** Quality of sequence reads was checked to obtain quality checks metrics using FastQC GUI. In the fastqc directory type;

~~~
$ ./fastqc
~~~

FastQC returns HTML report that consist of summaries and illustrations of base quality scores, GC, base and N content along with duplicated and adapter sequences (Fig.2).

**Figure 2.**
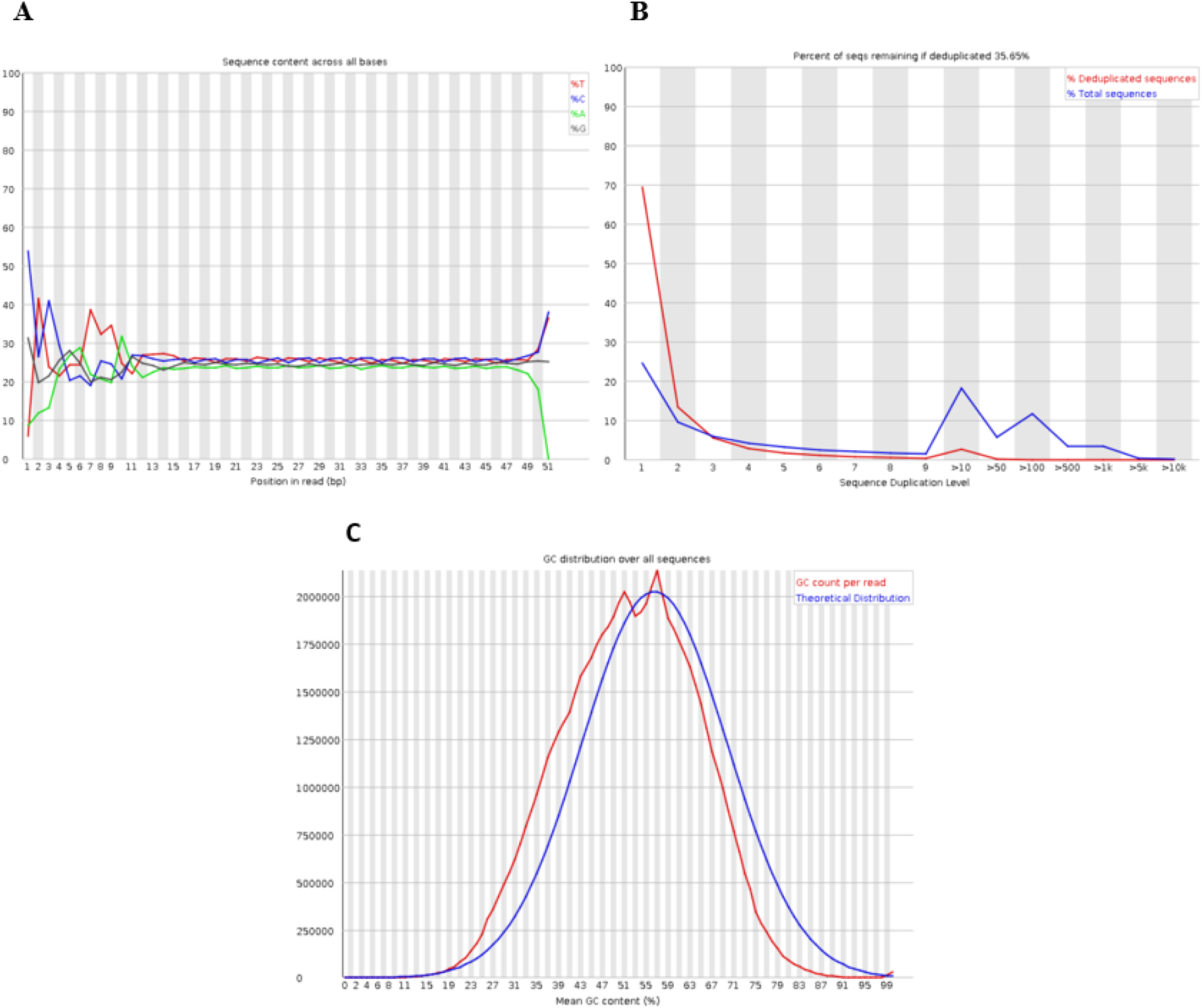
A graphical representation of FastQC report. **(A)**, per base sequence content; **(B)**, sequence duplication levels; **(C)**, per sequence GC content.

**2.** Low quality sequences were removed using Trimmomatic, applied separately on each pair of files.

~~~
$ java –jar /path to/Trimmomatic-0.36.jar PE –threads 3 fastq1.gz fastq2.gz trimmed1.fq untrimmed1.fq trimmed2.fq untrimmed2.fq SLIDINGWINDOW:4:20 MINLEN:2.
~~~

### Quasi-mapping and quantification of samples

**3.** Index the reference transcriptome FASTA file using the salmon indexer mode of wicked fast and accurate salmon tool as follows

~~~
$ salmon index –t transcripts.fa –i output_index_dir-name
~~~

When command was expedited, the output directory contained 10 files.

**4.** We then quantified set of paired-end reads separately in mapping-based mode against the index generated using salmon quant feature.

~~~
$ salmon quant –i output_index_dir-name -l A -1 trimmed1.fq -2 trimmed2.fq –validateMappings –o name_for_out-dir –numBootstraps 30
~~~

One may use the parameter, --gene_trans_map, giving gene(tab)transcript identifier file as input, for gene quantification. Directory containing 4 files and 3 sub-directories are created. The quant.sf file created contains quantification results in it (Table 1). Each file of it contained 47,159 entries.

**Table 1.** Format of the quantification results is displayed.

| Name | Length | Effective Length | TPM | NumReads |
| --- | --- | --- | --- | --- |
| NM_001285537.1 | 668 | 424.245 | 294.3024 | 2514.612 |
| NM_001285538.1 | 2507 | 2256.993 | 0.581311 | 26.424 |
| NM_001285539.1 | 786 | 538.318 | 0.184472 | 2 |
| NM_001285540.1 | 2391 | 2140.993 | 0.0693 | 2.988 |
| NM_001285544.1 | 639 | 396.707 | 10.01289 | 80 |
| .... | ..... | .... | .... | .... |

### Estimating transcript/gene expression abundances

**5.** For each of the sample, we built a matrices of counts and normalized values using abundance_estimates_to_matrix.pl script available in trinity toolkit.

~~~
$ perl abundance_estimates_to_matrix.pl --est_method salmon --gene_trans_map none --out_prefix name_for_output_files /path to/quant.sf /path to/quant.sf
~~~

This command generates three matrices files (i) TMM.expression.matrix (ii) counts.matrix (iii) TPM_not_cross_normalized.

### Number of expressed genes/transcripts counts

**6.** To infer approximate number of genes/transcripts that are properly mapped, quantified and expressed in our samples above some minimum threshold, we make use of TPM_not_cross_normalized matrix generated earlier and applied a perl script in trinity tool

~~~
$ perl count_matrix_features_given_MIN_TPM_threshold.pl TPM_not_cross_normalized_file | tee genes/trans_matrix.TPM.not_cross_norm.counts_by_min_TPM
~~~

(Table 2) represents the output which showed total 22,993 transcripts are expressed per TPM in any one of the subject samples.

**Table 2:**
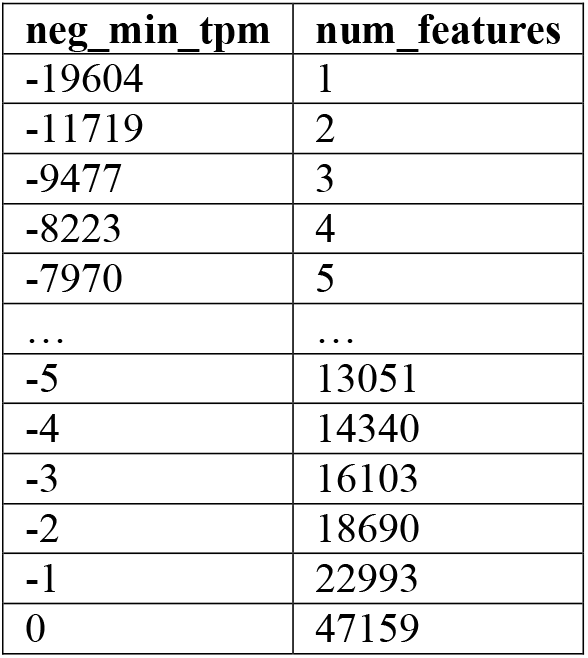
Output for transcript counts as a function of minimum TPM values.

| neg_min_tpm | num_features |
| --- | --- |
| -19604 | 1 |
| -11719 | 2 |
| -9477 | 3 |
| -8223 | 4 |
| -7970 | 5 |
| ... | ... |
| -5 | 13051 |
| -4 | 14340 |
| -3 | 16103 |
| -2 | 18690 |
| -1 | 22993 |
| 0 | 47159 |

**7.** The resulting output is then plotted on R using its plot function thus extrapolating numerous expressed features that have very low expression support (Fig. 3).

**Figure 3.**
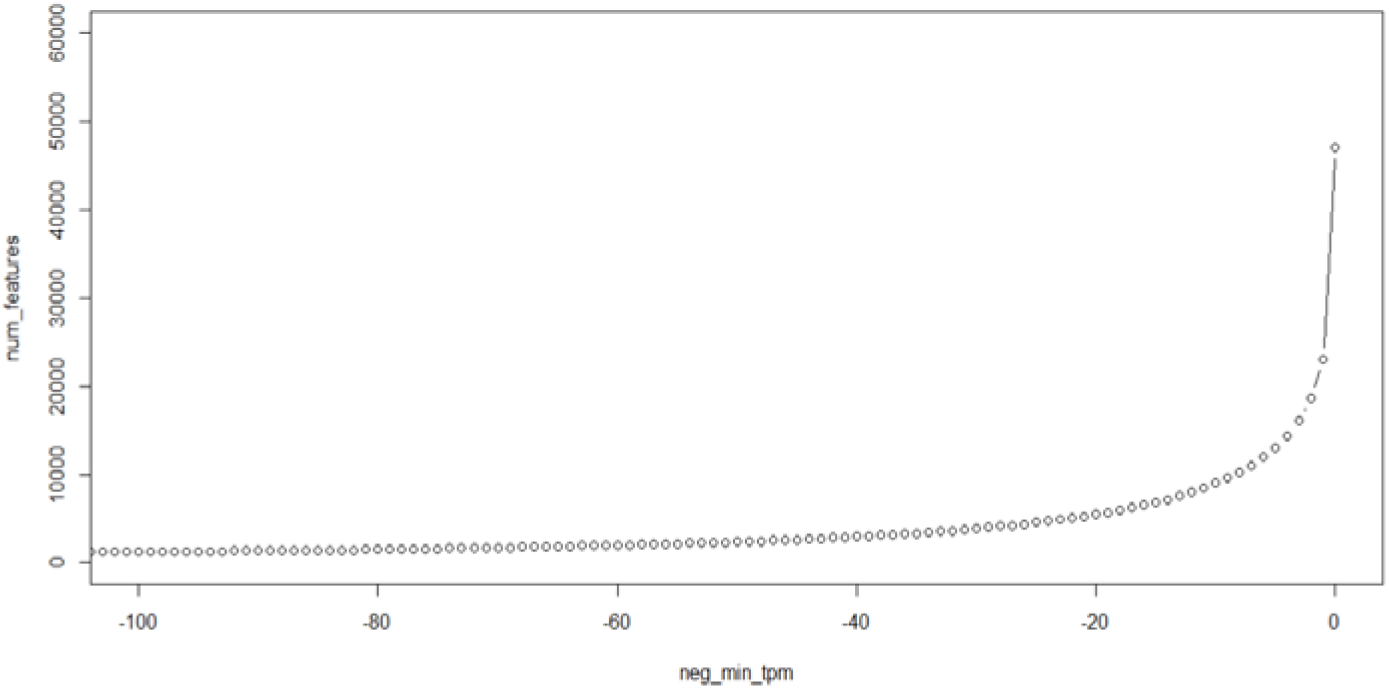
Number of genes/transcripts on x-axis are displayed against the TPM values of it on y-axis.

### Differential expression analysis

**8.** Pairwise comparison of both samples is performed on counts.matrix file which identified and clustered the DGE/transcripts according to most significant logFC, FDR, logCPM and P-values (Table 3).

**Table 3.** Layout of counts.matrix of differentially expressed isoforms.

|  | sampleA | sampleB | logFC | logCPM | PValue | FDR |
| --- | --- | --- | --- | --- | --- | --- |
| NM_001285727.1 | sampleA | sampleB | 15.82020742 | 6.922259796 | 1.69E-27 | 4.29E-23 |
| XM_018048107.1 | sampleA | sampleB | 15.6155561 | 6.71768419 | 6.95E-27 | 8.83E-23 |
| XM_005693226.3 | sampleA | sampleB | 14.31764944 | 5.420617384 | 5.43E-23 | 4.60E-19 |
| XM_005693223.3 | sampleA | sampleB | 13.99324758 | 5.096575019 | 5.08E-22 | 3.09E-18 |
| XM_018052275.1 | sampleA | sampleB | 13.96660044 | 5.069961231 | 6.09E-22 | 3.09E-18 |
| ... | ... | ... | ... | ... | ... | .... |

**9.** Furthermore, we extracted the differentially expressed features that have P-values at most 1e-3. This step relies on DESeq2, limma/voom and EdgeR Bioconductor packages and its dependencies. First command created a directory containing 4 files while the latter generated 12 files.

~~~
$ perl run_DE_analysis.pl --matrix counts.matrix --method edgeR --dispersion 0.1
$ perl analyze-diff-expr.pl --matrix TMM.EXPR.matrix -P 1e-3
~~~

Significant statistical inferences can be drawn from these values. For instace, the first value of logFC, 15.82 indicates that one of the sample is 2^15.82^ = 57848.82 times more expressed as compared to the other one, one doubling of the expression to the other one. The logFC versus normalized mean counts as MA plot, P-value vs. fold-changes for comparison between two conditions as volcano plot, genes having almost similar expression patterns are clustered togethere on heatmap and sample correlation matrix are illustrated in (Fig. 4).

**Figure 4.**
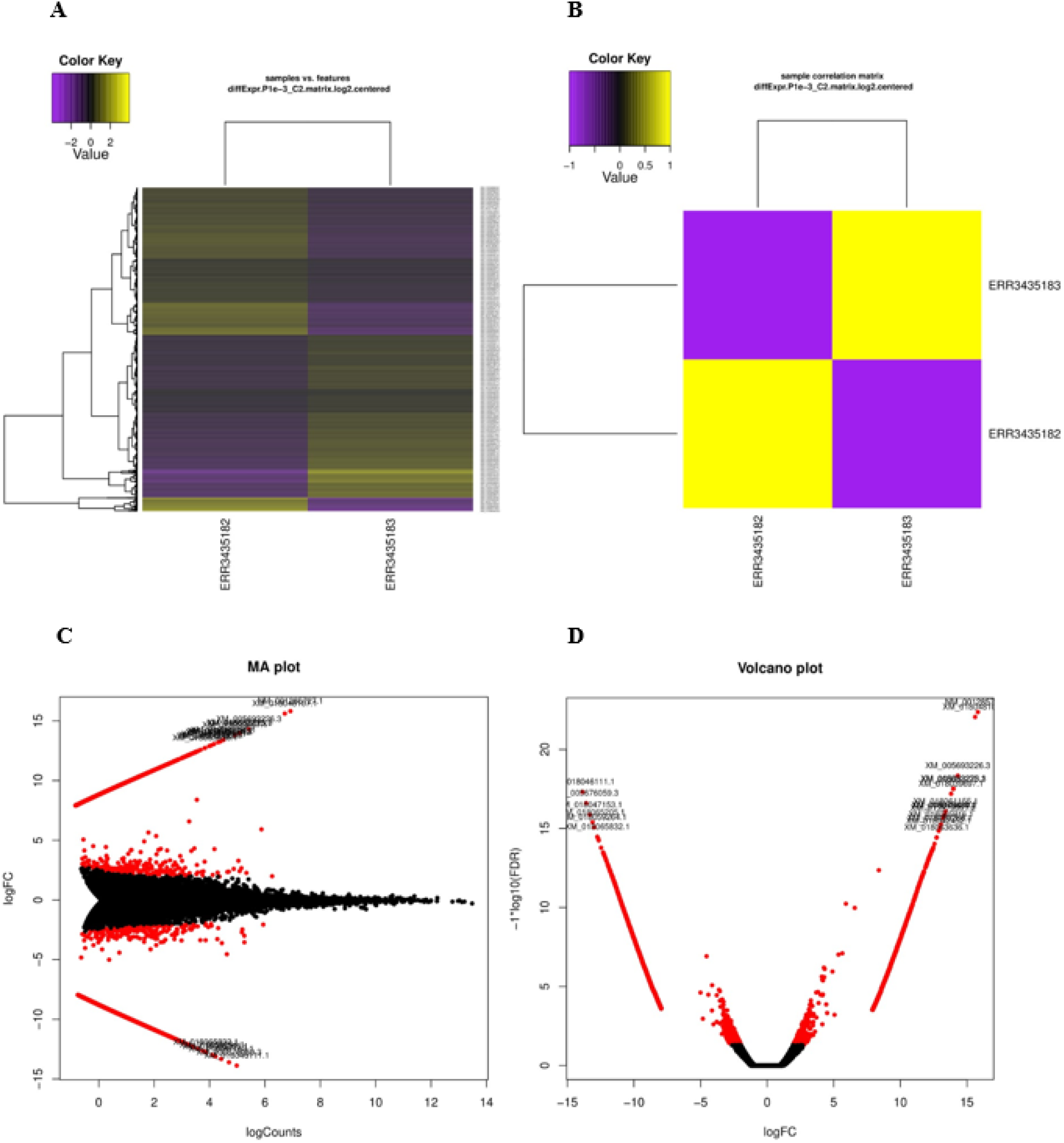
Results of DGE analysis. **(A)**, Results are plotted as MA plot**; (B),** Volcano plot; **(C)**, Heatmap; **(D),** Sample correlation matrix.

To describe this visual reference and to describe the expression matrix more precisely in terms of biological significance, some more detailed studies are still require and the pipeline to establish. We anticipate further researches to be conducted using this pipeline and onward integration of this workflow with functional annotations to the expression matrix pipeline in near future.

